# The potential function of forehead gland secretions in conflict resolution of the male Great Himalayan leaf-nosed bats

**DOI:** 10.1101/2020.09.03.281303

**Authors:** Chunmian Zhang, Congnan Sun, Jeffrey R. Lucas, Hao Gu, Jiang Feng, Tinglei Jiang

## Abstract

Chemical communication is an important aspect of social behavior in almost all animals. Here, we used gas chromatography-mass spectrometry (GC-MS) to detect the chemical composition, and behavioral tests to evaluate the potential function of forehead gland secretions between adult male Great Himalayan leaf-nosed bats, *Hipposideros armiger*. Our results showed that the concentrations of compounds and their categories differed significantly among individuals, and behavioral studies indicated that males are capable of utilizing the secretions for individual discrimination. Moreover, paired males that were incapable of gland protrusion showed more physical contact and longer contest duration compared to pairs in which both males could protrude the gland. In trials where only one male could protrude the gland, males with gland protrusion were more likely to win in contests. These findings provide the first behavioral evidence that chemical communication plays a vital role in conflict resolution in non-human mammals.

## Introduction

Contests over limited resources are ubiquitous in the animal kingdom (Bradbury & Vehrencamp, 2011). However, contests may increase injury risk and cost of energy, and may result in fatalities (Briffa & Elwood, 2004; Enquist & Leimar, 1990). Therefore, most animals tend to resolve conflict by exchanging information about their fighting ability, aggressive motivation or social status via multimodal signals including visual, acoustic or olfactory components before engagement in costly physical contests (Bradbury & Vehrencamp, 2011). Most of the studies on the effects of signals on conflict resolution have focused on visual and acoustic signals (Logue et al., 2010; Tibbetts & Lindsay, 2008); how olfactory signals affect conflict resolution is not well studied (Stritih & Kosi, 2017).

Chemical signals derive from complex blends consisting of several to hundreds of volatile or nonvolatile compounds expressed in glands, feces, urine and saliva. These signals mediate important social interactions in animals (Albone, 1984; Drea et al., 2013). Compared to visual and acoustic communication, chemical signals can be extraordinarily complex (Bushdid et al., 2014), in addition to maintaining signal integrity after a sender is absent, making them useful for decreasing the potential risks involved with physical contact between the senders and receivers (Alberts, 1992; Liebal et al., 2014). Indeed, chemical signals play a crucial role in animal contests and are used to defend territories, assess rivals and advertise multiple types of information such as individual identity, levels of threat/alarm, and social status (Burgener et al., 2009; Moore et al., 1997; Stritih & Kosi, 2017).

Chemical signals containing information about individual identity have been documented in various taxa, from insects (Lenoir et al., 1999), amphibians (Chouinard et al., 2013), fish (Sorensen, 2015), reptiles (Moreira et al., 2006), birds (Mardon et al., 2010), to mammals (Burgener et al., 2009). Individual recognition via chemical signals allows for stable interactions between known individuals. Individual recognition may also enable conspecifics to eavesdrop on contests between others from which the eavesdropper can gather information about the quality of both contestants. This information can then be used to challenge individuals with poor quality (Atema & Steinbach, 2007; Wyatt, 2014).

Chemical signals can also convey information about levels of threat/alarm, which could help to resolve conflicts during contests (Stritih & Kosi, 2017). For example, the chemical signals from the scent glands of cave crickets, *Troglophilus neglectus*, function to advertise the level of threat during agonistic interactions, which could prevent contest escalation and reduce conflict-related costs (Stritih & Kosi, 2017). So far, studies on chemical signals used during agonistic interactions have been reported in lobsters (Breithaupt & Atema, 2000), crayfish (Breithaupt & Eger, 2002), mice (Novotny et al., 1985) and hamsters (Rendon et al., 2016). In contrast, the role of chemical signals in mammals that function in conflict resolution resulting in de-escalation of interactions has been largely unexplored, especially in bats.

Bats provide a promising model to investigate the potential role of chemical communication in social behavior, because they have a variety of odor-producing glands on the chest, chin, face, genitals, shoulder, wing sac and subaxillary region which mediate their social activities between group members (Adams et al., 2018; Bradbury & Vehrencamp, 2011; Brooke & Decker, 1996; Faulkes et al., 2019; Rehorek et al., 2010; Scully et al., 2000; Voigt et al., 2008). Prior studies about bats’ chemical communication mainly focused on the information contained in their chemical signals, such as individual identity, colony identity, species identity, sex, age and mating status, and the potential role in individual or species discrimination, parent-offspring recognition, courtship and territorial scent-marking (Adams et al., 2018; Bloss et al., 2002; Bouchard, 2001; Caspers et al., 2008; Caspers et al., 2009; Faulkes et al., 2019; Gustin & McCracken, 1987; Safi & Kerth, 2003; Voigt et al., 2008). However, studies integrating chemical composition analysis and behavioral verification are largely unknown in bats, especially in the context of territorial conflicts.

The Great Himalayan leaf-nosed bat (*Hipposideros armiger*) is a highly gregarious species that usually roosts in caves, sharing day and night roosts among hundreds of individuals (Cheng & Lee, 2004). Our previous studies found that adult males in daily roosts usually maintain a minimum distance of 10-15 cm between individuals and defend their roosting territory using conspicuous agonistic displays such as broadband calls, wing flapping, boxing and wrestling (Sun et al., 2019). Our unpublished observations suggest that protrusion if the facial gland may also have a similar function (CM, pers. observ.). Adult male *H. armiger* possess a large forehead gland on their face that can emit a pungent, thick black secretion during contests (Figure 1a), an odor detectable by humans within a distance of 20 cm (CM, pers. observ.). The gland is only found in males (CM, pers. observ.).

**Figure 1.**
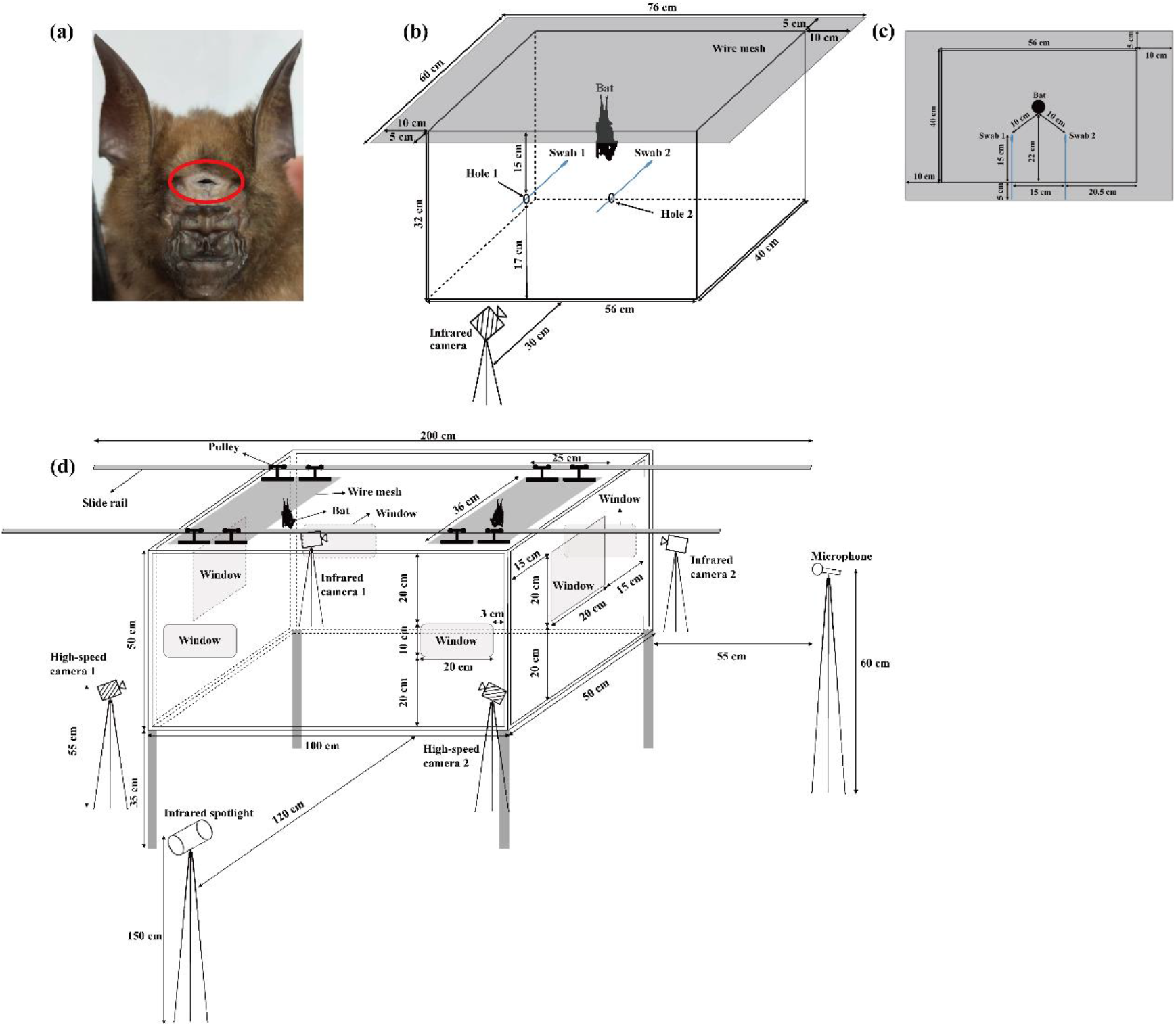
The forehead gland and experimental design. (a) Adult male *Hipposideros armiger*. Circle indicates the location of the forehead gland. (b) The habituation-discrimination test apparatus. Two swabs were used for presenting odor samples. In the habituation phase, the odor source of swab 1 was the habituation odor, and swab 2 was used as a blank control. In the discrimination phase, the odor source of swab 1 was the novel odor, and the odor source of swab 2 was the habituation odor. (c) Top-down view of the habituation-discrimination tests. The distance between the tested bat and each swab was 10 cm. Before the trial, the wire mesh could be rotated so that the tested bats were equidistant from the two odor sources. (d) Gland protrusion manipulation apparatus. The experiments were conducted in a 1.00 m × 0.50 m × 0.50 m box made of acrylic sheet plexiglass. The lid was removed. There are two windows on front and back to record the behavior of bats using infrared cameras, and one window on left and right to record the calls of bats using a microphone. Four pulleys slide steadily on two rails. Before the trial, the two bats roosted in the center of two pieces of mesh (grey area).

The aim of this study was to investigate the chemical composition and two potential functions of the forehead gland secretions we previously observed being used by *H. armiger* males during territorial conflicts. Since the forehead gland was often everted during roost territorial conflict, we hypothesized that chemical signals could play a crucial role in conflict resolution under these circumstances. Since individual recognition is a prerequisite for almost all social interactions including territory conflict (Tibbetts & Dale 2007), we first predicted that specific chemical compounds and their concentrations, as well as certain categories of compounds would vary among individuals thereby potentially providing information on individual identity. Second, we predicted that *H. armiger* males would have the ability to discriminate different individuals based on chemical compounds of forehead gland secretions. Finally, if the volatile odors of gland secretions contain chemical compounds conveying information relevant for conflict resolution, we predicted that the proportion of physical contact, contest duration, the frequency of aggressive displays and the proportion of aggressive displays would increase when gland protrusion was prevented in individuals during territorial conflicts. This latter prediction was tested in paired contests where either one or both of the paired bats had a sealed forehead gland that was incapable to secretion.

## Results

### Experiment 1: Chemical characterization of forehead gland secretions Scent profiles

We detected a total of 431 volatile compounds in 33 samples from 7 individuals (Mean ± SE = 8.76 ± 0.30 mg/sample; range: 5-13 mg). A total of 84 volatile compounds remained after filtering and these compounds were used for further statistical analyses (Table S1). Detected volatile compounds were classified into the following 16 categories: alkane, alcohol, aldehyde, ketone, ester, carboxylic acid, phenyl, ether, terpene, phenol, alkene, amine, peroxide, ammonium salt, nitrile and sulfone. Among them, alkane, alcohol, aldehyde, ketone and carboxylic acid accounted for 68% of the 84 compounds (Table 1). We tested for correlations between the five most common categories. There was a significant correlation in concentrations between alkane and aldehyde, alkane and ketone, alkane and carboxylic acid, and between alcohol and carboxylic acid (Table 2).

**Table 1.** Major categories of compounds identified from the forehead gland of 7 bats. The known function of each category is listed.

| Number | Category of compounds | Proportion in this study (%) | Function |
| --- | --- | --- | --- |
| 1 | Alkane | 19 | Recognition |
| 2 | Alcohol | 14 | Recognition/Threat/Alarm/Range marking |
| 3 | Aldehyde | 14 | Threat/Alarm |
| 4 | Ketone | 11 | Threat/Alarm |
| 5 | Carboxylic acid | 10 | Recognition/Threat/Alarm/Range marking |
| 6 | Others | 32 | Unknown |

Source: Alberts, 1992.

**Table 2.** Pearson correlation coefficients (*r*; below diagonal) and corresponding *p* values (above diagonal) of the major categories of compounds identified from the forehead gland of 7 bats.

| Category of compounds | Alkane | Alcohol | Aldehyde | Ketone | Carboxylic acid |
| --- | --- | --- | --- | --- | --- |
| Alkane |  | 0.199 | **0.011** | **0.002** | **0.044** |
| Alcohol | 0.552 |  | 0.371 | 0.194 | **0.023** |
| Aldehyde | **0.870** | 0.402 |  | 0.103 | 0.083 |
| Ketone | **0.943** | 0.557 | 0.665 |  | 0.092 |
| Carboxylic acid | **0.767** | **0.825** | 0.695 | 0.681 |  |

*Note:* Bold font indicates significant correlations ( $p < 0.05$ ).

#### Individual-specific scent profile

There were significant differences in chemical composition between individuals based on the relative peak area of each volatile compound (ANOSIM: Global *R* = 0.183, *p* = 0.008; Figure 2a). Based on the average of the relative peak area of each category of compounds, there were also significant differences in categories of compounds between different individuals (ANOSIM: Global *R* = 0.196, *p* = 0.004; Figure 2b). A descriptive comparison of absence or presence of volatile compounds in individual forehead gland was shown in Table 3. Individual A had four specific compounds that were not detected in other individuals; individual C had two specific compounds; individuals E and F had one specific compound each.

**Figure 2.**
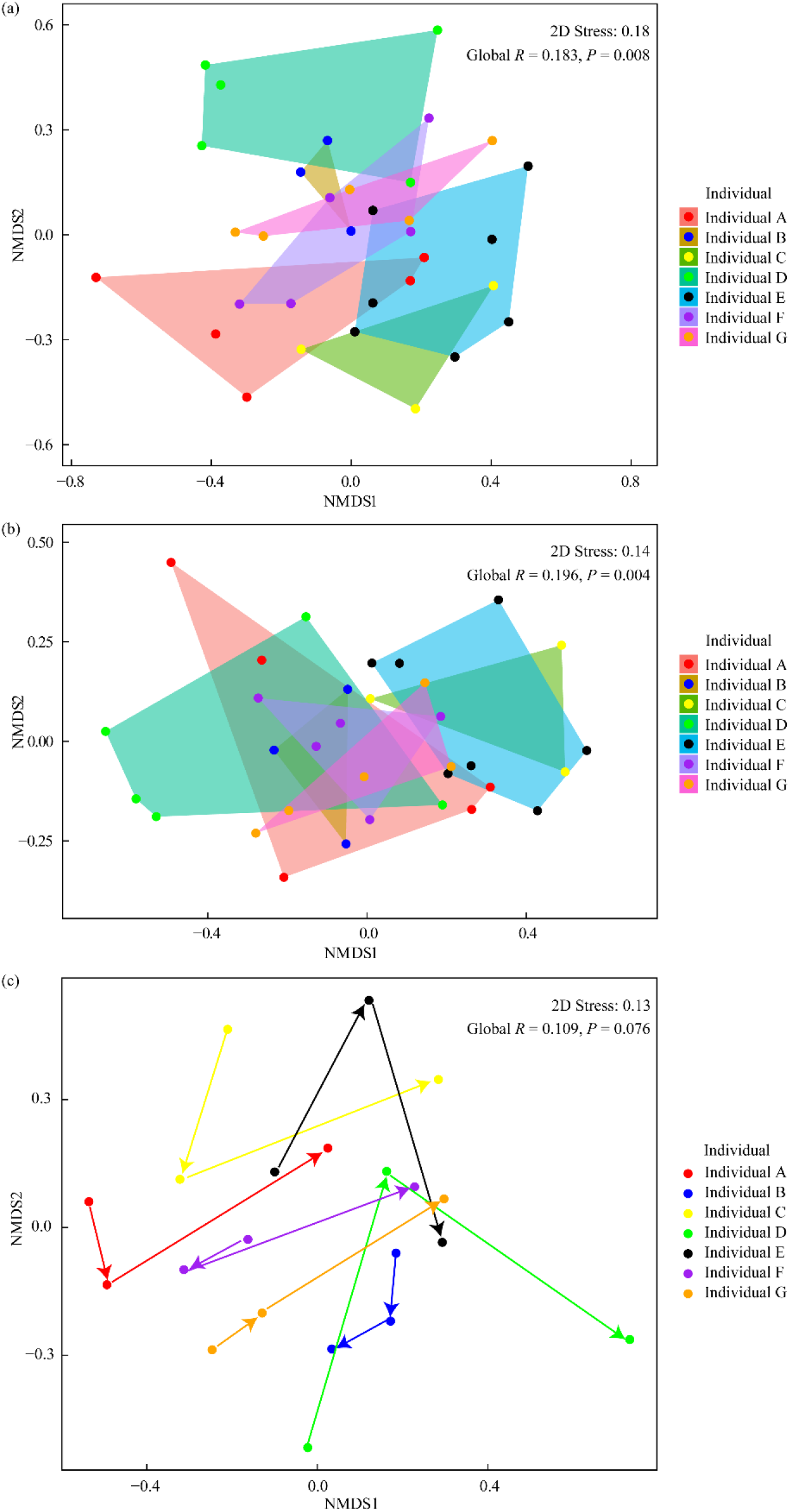
Non-metric multidimensional scaling plot showing (a) the similarity in chemical composition of 33 samples from 7 individuals, (b) chemical similarity of major categories of compounds identified from the forehead gland of 7 bats and (c) changes in the scent profiles of 7 individuals over a period of 3 months. Arrows indicate the sequence of sampling from the first to the last. Each colour represents one individual. Nearby samples have a similar scent and distant samples have a dissimilar scent. Axes are dimensionless.

**Table 3.** Description of the number of compounds in individual *Hipposideros armiger* gland secretion with a minimum of three repeat samples.

| Individual | Mean number of compounds per individual ± SE | Number of compounds occurring in all samples from the individual | Individually specific compounds in samples | N |
| --- | --- | --- | --- | --- |
| Individual A | 65 ± 2.4 | 38 | 4 | 5 |
| Individual B | 65 ± 1.5 | 50 | 0 | 3 |
| Individual C | 66 ± 1.2 | 49 | 2 | 3 |
| Individual D | 59 ± 3.4 | 30 | 0 | 5 |
| Individual E | 66 ± 1.3 | 39 | 1 | 7 |
| Individual F | 64 ± 2.9 | 41 | 1 | 5 |
| Individual G | 60 ± 3.2 | 38 | 0 | 5 |

*Note:* Letter ‘N’ indicates the number of samples from each individual.

Temporal patterns in chemical composition from the 7 individuals were tested using 3 samples per individual collected over a 3-month period (Figure 2c). For each individual, the chemical composition of the glandular secretions didn’t change significantly over time (ANOSIM: Global *R* = 0.109, *p* = 0.08; Figure 2c). Therefore, in this study, we removed the effect of different sampling dates on chemical composition.

### Experiment 2: Scent discrimination

We tested 12 males for their ability to discriminate the odors of glandular secretions from different individuals in habituation-dishabituation tests. In the habituation phase, the sniffing duration for the tested bats decreased across the four habituation trials. Pairwise comparisons showed that the sniffing duration decreased significantly from the first to the third presentation, and from the first to the fourth presentation of the habituation odor (Figure 3). These results indicate habituation to the odors. In the discrimination phase, the sniffing duration of the novel odor was significantly longer than the sniffing duration of the habituation odor (paired *t* test: *t*11 = 9.782, *p* < 0.001; Figure 3), which indicated a dishabituation response. These results suggested that bats could discriminate individual differences in odors of forehead gland secretions between different individuals.

**Figure 3.**
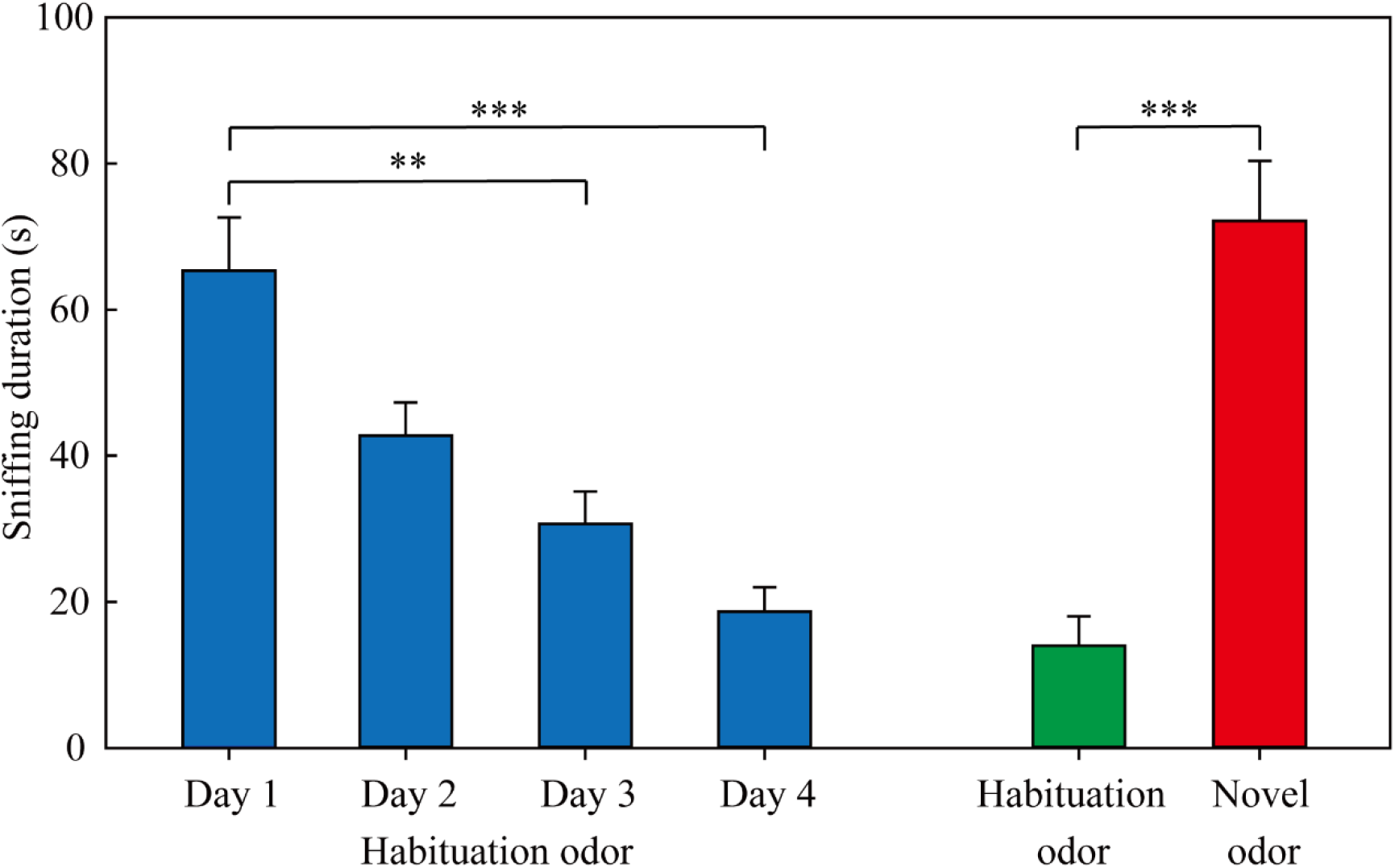
Sniffing duration of gland odors by *Hipposideros armiger* in the habituation-discrimination tests. Blue columns = habituation trials; green column = discrimination trials with habituation odor; red column = discrimination trials with the novel odor. Asterisks indicate significant differences (** *p* < 0.01; *** *p* < 0.001).

### Experiment 3: Effects of chemical signals on agonistic interactions

No significant differences were found in the contest duration between Yunnan population and Guizhou population (independent sample *t* test: *t*_18_ = - 1.148, *p* = 0.266). No significant differences were found in the proportion of bats using each behavioral pattern (i.e., wing flapping, boxing and wrestling) between Yunnan and Guizhou population (Pearson chi-square test: χ^2^ = 0.633, *df* = 2, *p* = 0.729). Therefore, we removed the effects of population on contest duration and agonistic displays.

For each pair condition, there were no significant differences in the contest duration (ANOVA: *F*_2,17_ = 1.946-2.906, *p* = 0.082-0.276; Figure S1a), frequency of aggressive displays (ANOVA: *F*_2,17_ = 0.672-1.832, *p* = 0.190-0.524; Figure S1b) or proportion of aggressive displays (Kruskal–Wallis test: *H*_2_ = 0.308-5.039, *p* = 0.081-0.857; Figure S1c, d, e) between the 3 vocalization categories (i.e., vocalization by both opponents, vocalization only by one opponent and no vocalization by both opponents). Therefore, we removed the effects of vocalization on contest duration and agonistic displays in this study.

Compared to the control condition, preventing gland protrusion significantly affected the levels of escalation expressed by both contestants, as well as the contest duration and fight outcomes (Figure 4). There were significant differences in proportion of physical contact among the 3 pair conditions (Pearson chi-square test: χ^2^ = 6.933, *df* = 2, *p* = 0.031; Figure 4a). Although there were no significant differences in proportion of physical contact between trials with both protruding males versus mixed pairs (Pearson chi-square test: χ^2^ = 2.849, *df* = 1, *p* = 0.091; Figure 4a), and between mixed pairs and pairs where both bats had no gland protrusion (Pearson chi-square test: χ^2^ = 0.784, *df* = 1, *p* = 0.376; Figure 4a), there was a significant difference in the proportion of physical contact between protruding pairs and unprotruding pairs (Pearson chi-square test: χ^2^ = 6.144, *df* = 1, *p* = 0.013; Figure 4a).

**Figure 4.**
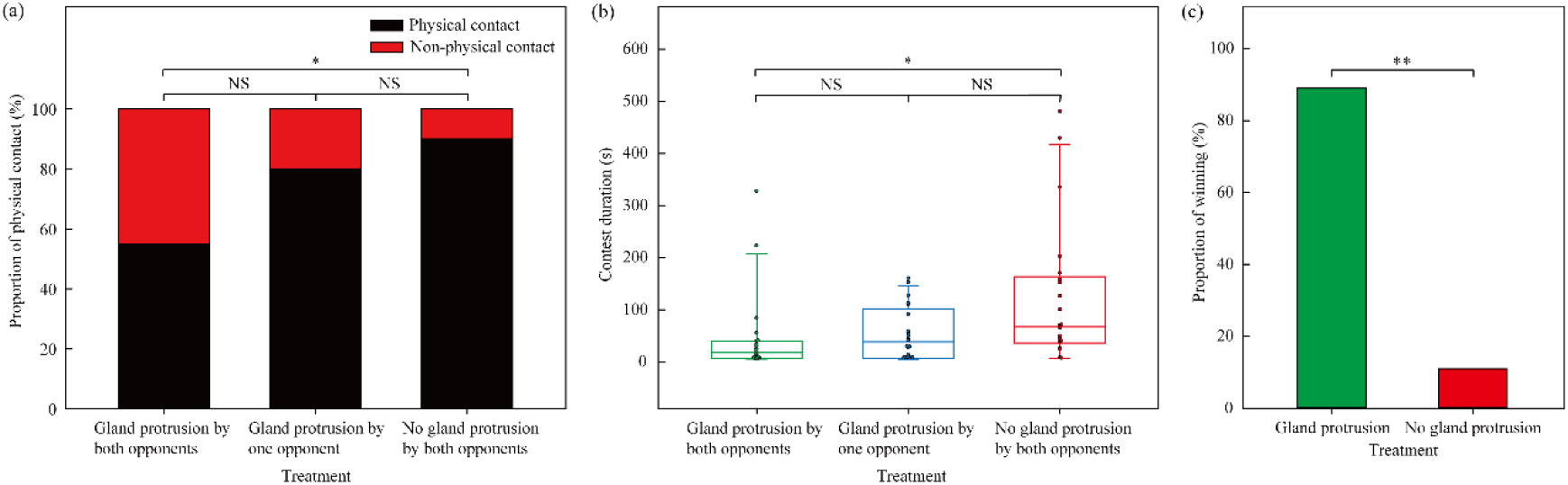
Effects of gland manipulation trials. (a) The proportion of physical contact in different treatments, (b) the contest duration in different treatments, and (c) the proportion of winning in a mixed pair (i.e. one with a functioning gland and the other with a glued gland). Asterisks indicate significant differences (* *p* < 0.05; ** *p* < 0.01), NS indicate no significant difference (*p* > 0.05).

There were significant differences in contest duration among the 3 pair categories (ANOVA: *F*_2,57_ = 4.64, *p* = 0.014; Figure 4b). Tukey’s multiple-comparison tests showed a pattern similar to that seen with the proportion of contests with physical contact: the only significant different between pair categories was for pairs with two protruding males compared to pairs where both males had no gland protrusion (Figure 4b).

In interactions between mixed pairs, we could determine a clear winner and loser in 18 of the 20 interactions. Among the 18 interactions, 16 (89%) were won by the individuals with gland protrusion, and 2 (11%) were won by the individuals with a glued gland. As a result, the individuals with gland protrusion were significantly more likely to win during agonistic interactions (binomial test: *p* = 0.001; Figure 4c).

No significant differences were found in the frequency of aggressive displays among the 3 pair categories (ANOVA: *F*_2,57_ = 1.946, *p* = 0.152; Figure S2a). There were also no significant differences in the proportion of wing flapping, proportion of boxing and proportion of wrestling among 3 pair conditions (Kruskal–Wallis test: *H*2 = 0.567-5.940, *p* = 0.051-1.133; Figure S2b). In the mixed pairs, no significant difference was found in the frequency of aggressive displays between the individual with gland protrusion and the individual with no gland protrusion (paired *t* test: *t*_19_ = - 0.418, *p* = 0.681; Figure S2c). Moreover, there were no significant differences in the proportion of aggressive displays between the individual with gland protrusion and the individual with no gland protrusion (Wilcoxon signed-ranks test: wing flapping: *Z* = - 0.973, *N* = 20, *p* = 0.331; boxing: *Z* = - 0.910, *N* = 20, *p* = 0.363; wrestling: *Z* = - 1.376, *N* = 20, *p* = 0.169; Figure S2d).

## Discussion

In this study, we detected 84 volatile compounds falling into 5 major categories of compounds: alkane, alcohol, aldehyde, ketone and carboxylic acid. Among them, alkanes have been shown to serve the function of recognition, while aldehydes and ketones have been shown to serve the function of threat (Alberts, 1992). We also found that there were significant correlations in concentrations between alkane (recognition function) and aldehyde and ketone (threat function). These results supported our first prediction that the volatile odors of gland secretions contain chemical compounds that potentially serve recognition and threat functions. Moreover, there were significant differences in the concentrations and categories of compounds between different individuals, and all tested bats could discriminate between different individuals based on the odors of forehead gland secretions in the habituation-dishabituation trials, supporting our second prediction that chemical compounds of gland secretions convey information on individual identity, and can be used for individual discrimination. Finally, we found that the proportion of physical contact and contest duration significantly increased when gland protrusion was prevented in both opponents, supporting our third prediction that the presence of gland secretions will impact contest resolution. Based on the empirical support of these predictions, our results strongly confirmed that chemical signals of forehead gland secretions in *H. armiger* could help to resolve conflicts during agonistic interactions over roost territories.

### Multiple information

We found that there were significant differences in the concentrations of compounds and their categories between different individuals. These results suggest that the odors of forehead gland secretions of *H. armiger* convey information about individual identity. Similar results have been found in the Bechstein’s bat *Myotis bechsteinii* (Safi & Kerth, 2003), greater spear-nosed bats *Phyllostomus hastatus* (Adams et al., 2018), and male greater sac-winged bats *Saccopteryx bilineata* (Caspers et al., 2008). Selection assuring the ability of territory owners to signal their individual identity is especially crucial in male *H. armiger*, because they live in a large and complex social system and are involved in agonistic interactions on a daily basis. The differences in chemical signals among individuals were useful for individual discrimination, and thus may facilitate social interactions within and between colonies (Tibbetts & Dale, 2007). Moreover, we found that alkane, alcohol and carboxylic acid accounted for 43% in the forehead gland secretions of *H. armiger*. Prior studies show that the presence of functional groups of carboxyl, hydroxyl and aromatic rings in compounds could increase chemical stability and durability, and they serve the function of recognition and range marking because these compounds last a long time and are resistant to degradation from water and heat (Alberts, 1992; Bradbury & Vehrencamp, 2011). Therefore, our results suggested that the gland secretions of *H. armiger* contained chemical compounds that facilitate individual recognition.

We found that aldehyde and ketone accounted for 25% in the forehead gland secretions of *H. armiger*. Prior studies showed that the compounds with carbonyl functional groups are more soluble in water than those with carboxyl or hydroxyl functional groups. This characteristic promotes rapid volatilization and potentially serves a function of threat or alarm (Alberts, 1992; Bradbury & Vehrencamp, 2011). Therefore, our results implied that the gland secretions of *H. armiger* contained chemicals compounds serving a threat function. Moreover, we also found that there was a significant correlation in concentrations between chemicals that generally have recognition and threat/aggression functions, while there was no significant correlation in concentrations between chemicals with the same function. These results suggested that the functions of individual discrimination and mutual threat between two rivals may be coupled during agonistic interactions in *H. armiger*. These results also implied that the chemical compounds of forehead gland secretions convey multiple sources of information (i.e., recognition and threat).

Why do chemical signals of *H. armiger* encode multiple information? One possible explanation is that different compounds of gland secretion contain different information on the signalers’ attributes, and thus permits receivers to more comprehensively assess signalers’ fighting ability or resource holding potential (RHP). *Hipposideros armiger* usually roost in a dark cave, increasing the relevance of non-visual signaling. Intruders and defenders may assess fighting ability or RHP based on information of chemical signals, such as body size, aggressive motivation, offensive and defensive threat, and thus allow them to decide to withdraw or escalate in a contest. However, obviously only further experimental examination will help in answering how *H. armiger* use their chemical signals to assess fighting ability or RHP during the process of conflict resolution.

### Individual discrimination via chemical signals

Significant variation in sniffing duration to a habituation odor and to a novel odor provided behavioral evidence that *H. armiger* could discriminate the odors of forehead gland secretions from different individuals. Similar results have been found in Antarctic seabird *Pachiptila desolata* (Bonadonna & Nevitt, 2004), mice *Mus domesticus* (Hurst et al., 2001) and Belding’s ground squirrels *Spermophilus beldingi* (Mateo, 2006). The forehead gland secretions of *H. armiger* were a chemically complex substance consisting of a variety of volatile compounds. The individual-specific odor profile was determined by both the presence of individual-specific compounds and individual variation in the relative concentration of compounds. Multiple sources of individual variation have been described in several other mammals (Burgener et al., 2009; Hagey & MacDonald, 2003; Safi & Kerth, 2003; Smith et al., 2001). *Hipposideros armiger* has a harem-polygynous mating system, and a harem includes one territorial male and several females (Yang, 2011). Because harem males defend their females or a territory against other males, the ability of harem males to discriminate opponents based on chemical signals could mitigate energy expenditure through decreased confrontation with males with high competitive ability.

### Chemical signals for conflict resolution in territory defense

In agonistic interactions, males with sealed forehead glands were likely to lose to males with functioning glands. These results suggest that chemical signals of *H. armiger* may serve threat and/or alarm functions. We also found that interactions between males that could not protrude their glands were characterized by a higher proportion of physical contact (high aggression interactions) and longer contest duration than interactions between males that were capable of protruding their glands. This is an important result because it shows that the increased probability of losing in males with non-functioning glands is not a result of reduced vigor caused by gland manipulation. Indeed, we found the opposite pattern – the lack of a functioning gland actually increased the duration and intensity of the interaction. These results indicate that chemical signals of *H. armiger* may function as an offensive and defensive threat, which can inhibit contest escalation and reduce contest duration. Similar data have been found in cave crickets, *Troglophilus neglectus*, where the frequency of fighting and contest duration significantly increased after chemical signals were prevented in both contestants (Stritih & Kosi, 2017). Because the level of aggression and contest duration are strongly associated with the cost of conflict in terms of the energy expenditure, injury or risk of predation (Briffa, 2015; Briffa & Sneddon, 2007), we therefore suggest that chemical signals of *H. armiger* can mitigate the costs of conflict during territory defense.

We found no significant differences in the frequency and proportion of aggressive displays among 3 pair conditions and between the individual with gland protrusion and the individual without gland protrusion. These results indicated that chemical signals of *H. armiger* may have no influence on the displays of aggressive behavior. One possible interpretation is that the frequency and proportion of aggressive displays in a dyadic agonistic interaction of male *H. armiger* may depend on the RHP asymmetry between opponents, i.e., the total displays of aggressive behavior between size-matched opponents are similar. For example, in mantis shrimp *Neogonodactylus bredini*, there was no significant relationship between total number of strikes and mean opponent mass in size-matched contests (Green & Patek, 2018).

### Social vocalization and agonistic interactions

Acoustic communication can serve the function of reducing the cost of agonistic interactions (Logue et al., 2010). Acoustic signals can also convey the information of individual identity and aggressive motivation, which can make contestants decide whether to continue or cease fighting (Bradbury & Vehrencamp, 2011). Similar results have been obtained in male Seba’s short-tailed fruit bats *Carollia perspicillata* (Fernandez et al., 2014), Asian particolored bats *Vespertilio sinensis* (Zhao et al., 2018), and Indian False Vampire bats *Megaderma lyra* (Bastian & Schmidt, 2008). *Hipposideros armiger* use agonistic displays to defend their roosting territory, accompanied by vocal signals (Sun et al., 2019). Given these previous results, it is interesting that the presence or absence of vocal signals had no effect on contest duration or on the rate of agonistic displays. Moreover, our previous study showed that social vocalization during territory defense in *H. armiger* only encodes information on individual identity and emotional state (Sun et al., 2018), implying that the signals may not help to resolve conflicts. Thus, these results suggested that social vocalization of *H. armiger* may not significantly affect agonistic interactions for territory conflict. However, additional multimodal playback experiments including acoustic and chemical signals are needed to demonstrate the relative importance of vocalizations and odors in conflict resolution of *H. armiger* during territory defense.

### Conclusions

In summary, our study demonstrates that the chemical signals of forehead gland secretions of male *H. armiger* convey multiple sources of information about individual identity and threat/alarm. Further behavioral testing showed that chemical signals could decrease the proportion of physical contact and contest duration, suggesting that the chemical signals may function as an offensive or defensive threat. Our results from the chemical composition analysis and behavioral trials strongly confirmed that chemical signals of in *H. armiger* could help to resolve conflicts during agonistic interactions for roost territories. To our knowledge, our study provided the first behavioral evidence that chemical communication plays a vital role in conflict resolution in a non-human mammal. A limitation of this study is that it is difficult to determine if the chemical signals of *H. armiger* are associated with body size, fighting ability, or aggressive motivation based on our present data. Nonetheless, our study highlights the importance of chemical signals in conflict resolution. Further studies will need to determine what compounds play a part in individual recognition and conflict resolution, and determine the relative importance of chemical and acoustic signals during conflicts.

## Materials and methods

### Experiment 1: Identification of volatile compounds in forehead gland secretions Animals and housing

In April 2018, seventeen adult males of *H. armiger* were caught in a mist net from the Shiyan cave in Chongyi, Jiangxi Province, China. Males were classified as adults (> 1 year old) if they had a sealed epiphyseal gap, brown fur, and worn canine cusps (Cheng & Lee, 2002). All bats were kept in individual cotton bags and brought to the laboratory. Bats were housed in a husbandry room (6.5 m long × 5.5 m wide × 2.1 m high). The room was maintained at a temperature around 23°C, a relative humidity around 60% and a 12-hours dark/light cycle. All bats were fed with ad libitum freshwater and larvae of *Zophobas morio* enriched with vitamins and minerals. All bats were marked with metal rings (4.2 mm; Porzana Ltd, East Sussex, UK) on the forearm to identify individuals. The bands did not affect the normal behavior of the bats (Sun et al., 2018).

#### Scent collection

The forehead gland forms a deep pocket above the nose which can be everted when palpated (Figure 1a). We extruded the black secretion by squeezing the area around the forehead gland, and transferred it into a 20 ml glass headspace vial with a PTFE-lined septum using pre-sterilized forceps. We collected all samples between 19:00 and 19:30. To exclude the effect of potential contaminants, one blank sample was collected during each sampling period.

We collected 33 samples from 7 individuals (Mean ±SE = 4.7 ± 0.5 samples/individual; range: 3-7 samples/individual). We attempted to collect seven sequential samples per individual to test for within-individual variation in secretion properties over time. Sampling was attempted every 15 days for each of the bats because it typically took about 15 days for the gland to replenish its contents after palpation (CM, pers. observ.). However, we were unable to collect a usable sample from each individual every 15 days due to a minimal secretion or a complete lack of extruded secretion for some of the sampling periods.

#### Chemical compounds analysis

Before all samples were analysed, 10 μL of 2-Octanol (10 mg/L stock in dH2O) were added as an internal standard. The mixed sample was heated for 15 min at 60°C. Then each sample was extracted for 30 min in headspace solid-phase microextraction (SPME) using 50/30μm DVB/CAR/PDMS SPME fiber coating. After the volatile compounds were extracted, they were desorbed from the SPME fiber coating when immediately inserted at 250°C into the injector port.

The gas chromatography–mass spectrometry (GC–MS) analyses were performed with an Agilent 7890 gas chromatograph system linked to an Agilent 5977 mass spectrometer with the EI ion source (70 eV). The system utilized a DB-Wax capillary column (30 m × 250 μm inner diameter and 0.25 μm film thickness; Agilent, USA). A 1 μL sample was injected into in a 1:1 split mode. GC–MS analyses were performed with helium (at 1mL min^-1^) as the carrier gas, the front inlet septum purge flow was 3 mL min^-1^. The initial temperature was kept at 40°C for 4 min, then increased to 245°C at a rate of 5°C min^-1^, and then kept at 245°C for 5 min. The front injection, transfer line, and ion source temperature was 250, 260, and 230°C, respectively. The mass spectrometer was conducted in the full-scan mode with the m/z range of 20–500, solvent delay of 0 min.

Chroma TOF 4.3X software of LECO Corporation and National Institute of Standards and Technology (NIST) database were used to measure raw peaks, data baseline filtering and calibration of the baseline, peak alignment, deconvolution analysis, peak identification, integration and spectrum match of the peak area (Kind et al., 2009). Volatiles with less than 80% similarity compared to compounds in the NIST library and relative peak areas less than 0.1% were excluded from further statistical analyses. The relative peak area was calculated by dividing peak area of each compound by that of the peak area of the internal standard (IS) in the same analytical run. We ran a blank sample as a control to determine compounds which were derived from gland secretions of the bats and therefore considered to be endogenous. Compounds were assumed to be contaminants or exogenous compounds if they were in similar or higher concentrations in the blank sample than in the gland secretion sample. To avoid false positive compounds, only compounds detected in at least half of the samples from each individual were used for further analysis.

### Experiment 2: Individual discrimination

#### Scent collection

We collected 60 samples from 12 individuals (5 samples/individual) for the habituation phase and 12 samples from 12 individuals (1 sample/individual) for the discrimination phase. As with samples used to test for individual variation, replicate samples for the habituation-discrimination tests were collected at intervals of 15 days. We were able to collect a full set of samples from each bat for this part of the experiment. After collection, we weighed the secretions to within ± 0.001 g (AR2140, Ohaus International Trading Co. Ltd, China). All samples were stored at - 80°C until used.

#### Habituation-discrimination tests

We used habituation–discrimination tests to determine whether the bats could distinguish individual differences in the odor of the gland secretions (Halpin, 1986; Schultze-Westrum, 1969). In the habituation phase of this test, the subject was presented with two swabs once a day for four days. One odor was from a single individual (‘habituation odor’) and the other odor was from an unscented swab. The unscented swab was used as a control to verify that the tested bats habituated to the odor instead of the swab. The sniffing duration of the scented swab was significantly longer than the unscented swab in all habituation trials (paired *t* test: all *P* < 0.001). The bats were considered habituated to the odor if the sniffing duration decreased significantly across the four habituation trials. The data on sniffing duration of unscented swabs are not presented.

The discrimination phase was conducted on the fifth day. This phase involved presenting the bat with two odors, one was the ‘habituation odor’ and the second was an odor from another individual (‘novel odor’). This phase was used to determine whether the tested subject could discriminate between the two odors. If the tested subject showed a stronger response to the novel odor compared to the habituation odor, we assumed that the tested subject could discriminate the two odors. The side on which each sample was placed was randomly selected.

We performed all habituation-discrimination tests in a scentless plastic box without a plastic top (0.56 m long × 0.40 m wide × 0.32 m high; Figure 1b) in a 4.5 m long × 2.4 m wide × 2.2 m high room. All tests were conducted between 19:00 and 22:00. The top of the box was covered with a wire mesh (0.76 m long × 0.60 m wide), which enabled the bats to hang from the roof. The two odor sources were presented through 1 cm diameter holes in the side of the box. The holes were placed 15 cm from the wire mesh, which was the average distance from a bat’s head to toe. The distance between the two holes was also 15 cm. The wire mesh on the roof could be rotated so that the bats were facing directly toward and were equidistant from the position of the odor sources.

For the habituation phase, the bats were released into the center of the experimental set-up at least 5 min before the two swabs were presented. Before the trial, we placed vials containing the gland secretions on ice until the secretions unfroze completely (usually 10-20 min). After the bats became motionless, we placed 5 mg of gland secretions on a swab (20 cm in length and 2.8 mm in diameter) and inserted two swabs into two pieces of foam to stabilize them. We then moved both swabs slowly and simultaneously toward the bat a constant rate. The distance between each swab and the bat was 10 cm (Figure 1c), mimicking natural conditions. Considering the rapid volatilization of volatile compounds in gland secretions, the duration of each experiment was 10 min. We recorded the behavior of the bats for 10 min via an infrared camera (FDR-AX60; Sony Corp., Tokyo, Japan), which was placed 0.3 m in front of the box. Sniffing was defined as the bats moving their noses towards the swab up to a distance of ≤ 1 cm, and spending at least 1 s on it at a time. Each odor sample was used only once. If a swab was licked or dislodged, the data collected from that pair of swabs was ceased. After each trial, we cleaned the plastic box using 75% ethyl alcohol to remove volatile compounds, and we ventilated the room by opening a set of windows.

### Experiment 3: Manipulation of forehead gland protrusion during agonistic encounters Animals and housing

To test the role of chemical signals in conflict resolution, we caught 120 adult males from 2 localities in southern China, including individuals from Yunnan (96 males) and Guizhou (24 males) in July-August 2019. We captured at most 16 adult males at a time, and captured bats every one or two days. There was a turnover of bats every one or two days. Captured bats were housed for at least 24 h in individual cage (0.5 m long ×0.5 m wide ×0.5 m high) in a makeshift laboratory (6.0 m long ×3.4 m wide × 2.9 m high) before the experiment started. The room was maintained at a relative humidity around 65% and a temperature between 20 and 25 °C. Bats were given ad libitum freshwater and larvae of *Zophobas morio* enriched with vitamins and minerals. Each bat was used only once.

#### Morphological measurements

Body mass of each individual was measured using an electronic balance (± 0.01 g; DH-I2000, Diheng Ltd., Shenzhen, China), and we measured the length of the right forearm of each individual using an electronic vernier caliper (± 0.01 mm; 111-101V-10G, Guanglu Ltd., Shenzhen, China) before the trials. We measured the body mass and forearm length of each individual three times, and their averages were used for further analysis.

#### Staged agonistic interactions for territory

We conducted agonistic interactions between pairs of male *H. armiger* in a box made of acrylic sheet plexiglass without a lid (1.00 m long × 0.50 m wide × 0.50 m high; Figure 1d). The box was placed on two benches 0.35 m above the ground in a temporary laboratory (6.00 m long × 3.40 m wide × 2.90 m high). The temperature of the room was 20-25°C and the humidity was 50-70%. We placed two infrared high-speed cameras (Photonfocus MV1-D1312IE-240-CL-8) on two front windows to record the bats’ behavior and to detect the presence of gland protrusion. The two infrared high-speed cameras filmed behavior at 85 frames/s and the recordings were played back at 25 frames/s (Video S1). We placed two infrared cameras (FDR-AX60; Sony Corp., Tokyo, Japan) on two back windows to record the duration of any aggressive interaction. Two infrared high-speed cameras and two infrared cameras were mounted 0.55 m above the ground. We also set up an infrared spotlight (KTJ-GY-300W-42V; RockeTech Corp., Ltd., Hunan, China), which was mounted 1.20 m away from the box and 1.50 m above the ground, for sufficient illumination of the infrared high-speed cameras. We used an Avisoft UltraSoundGate116H (Avisoft Bioacoustics, Glienicke, Germany) consisting of a condenser ultrasound microphone (CM16/CMPA, Avisoft Bioacoustics, Glienicke, Germany) at 0.55 m from the right window of the box to record the bats’ vocalizations. The sampling frequency was at 250 kHz with 16 bit resolution. Pairs of males were selected randomly and placed individually in the center of the two pieces of wire mesh (0.36 m long × 0.25 m wide) on opposite ends of a slide rail (2.00 m long). Each piece of wire mesh was fixed by four pulleys to slide steadily on these two rails.

We removed the hair around the forehead gland before the trial to clearly detect the presence of a protrusion of the forehead gland and to avoid recapturing the tested bats. To exclude the effect of body size on the contest duration, intensity and outcome, we staged dyadic agonistic interactions between the paired males matched in body mass (0.9 < the ratio of body mass between males < 1.1; Sun et al., 2019) and in forearm length (the difference between males in forearm length /male-average length < 2% ± 1.2%; Green & Patek, 2015). We carried out trials where individuals either both were capable of gland protrusion, or neither was capable of gland protrusion, or one could and one could not protrude the gland. We eliminated the possibility of gland protrusion by gluing the facial cuticle over the glandular region, thereby completely preventing the forehead gland from protruding during the experiment. The glue was easily removed after the experiment. The adhesive had solidified before the onset of the experiment, so it no longer had any volatile odor. We used 60 pairs of males (20 pairs/group) in this experiment.

Before the trial, we randomly selected two males matched in body size and placed them in the center of the two pieces of wire mesh to allow them to acclimate to the box. After the two males remained motionless for at least 5 min (i.e., no body movements), we pulled the two pieces of wire mesh (approximately 0.5 m apart) attached to four pulleys slowly and simultaneously at a constant rate towards each other by a rope until they reached the center of the box; this simulated the situation of two bats invading each other’s territory. The distance between the two bats was approximately 15 cm. The process of pulling the wire mesh towards each other occasionally disturbed one of the bats causing it to fly off the mesh. We discarded such unsuccessful trials. The tested bats from unsuccessful trials were returned to the individual cages and were used in subsequent attempts. Agonistic interactions were terminated when we could determine a clear winner and loser within 15 min of the start of the trial. The trial was terminated if a winner could not be determined within the 15 min interval. We defined winner as the individual that remained in place after the loser had retreated. We defined loser as the individual that retreated after an agonistic interaction and didn’t display any aggressive behaviors after retreat for at least 20 s. Gland protrusion manipulation experiments were performed during the maximal agonistic interaction activity period (between 20:00 and 08:00). After the trial, we cleaned the box using 75% ethyl alcohol to exclude the potential effect of odor on the next trial, and the room was well ventilated to remove odor residues.

#### Behavioral analysis

We used QvodPlayer (Version 5.0.80, Shenzhen Qvod Technology Co., Ltd., Guangdong, China) to analyze video recordings from 60 dyadic agonistic interactions and to describe agonistic behavior. To compare the differences in contest duration and intensity among 3 trial conditions, we analyzed contest duration, proportion of physical fight, proportion of non-physical fight, frequency of aggressive displays and proportion of aggressive displays (for description of contest duration and behavioral parameters see Table 4; Clement et al., 2006; Sun et al., 2019).

**Table 4.** Description of contest duration and behavioral parameters of male *Hipposideros armiger*.

| Measurement | Description |
| --- | --- |
| Contest duration | The difference in time between the first wing flapping and the last wing flapping or boxing given by one of the opponents |
| Number of wing flapping | An up-and-down flapping cycle was recorded a wing flapping |
| Number of boxing | A boxing was recorded when the bat threw a punch with its wrist at opponent |
| Number of wrestling | A wrestling was recorded when two individuals held each other with their semi-extended wings or forearms when they fought |
| Physical fight | The fighting behavior of any physical contact during an aggressive interaction (i.e., boxing or wrestling) |
| Non-Physical fight | The fighting behavior of no physical contact during an aggressive interaction (i.e., wing flapping) |
| Frequency of aggressive displays | The total number of aggressive displays (i.e., wing flapping, boxing and wrestling) performed by each opponent divided by the contest duration |
| Proportion of aggressive displays | The number of wing flapping, boxing, wrestling performed by each opponent divided by the total number of aggressive displays, respectively |

#### Statistical analysis

To examine whether there were individual differences in chemical composition of gland secretions from different individuals, we calculated the relative peak area of the GC-MS peaks using the peak area of each compound divided by the peak area of the internal standard (IS). Based on the relative peak area, we calculated the Bray–Curtis similarity index between each pair of samples, and then used this to perform a non-metric multidimensional scaling (NMDS) ordination. This placed each sample in a two-dimensional space so that the relative distance between the samples matched their chemical similarity. ‘ Stress’ was used to measure ‘goodness of fit’, which assessed how well a particular configuration recreated the observed distance matrix associated with the data. We used the following criteria for stress results: stress < 0.05 showed an excellent representation in two dimensions; 0.05< stress < 0.1 was very good; 0.1< stress < 0.2 was good and stress > 0.2 showed a poor representation in two dimensions.

In this study, we determined the chemical composition of gland secretions from 7 individuals which contributed at least three samples. We computed a non-parametric analysis of similarities (ANOSIM) with 1,000 permutations on the basis of the Bray–Curtis similarity distance matrix of each individual. The ANOSIM test is a series of Mantel-type permutation or randomization procedures that do not require any assumptions about the distribution of data (Clarke & Warwick, 2001). We defined Global *R* as the difference in average rank dissimilarity within groups (here individuals) compared to between groups. Global *R* is 1 if all samples from the same group are different than samples from different groups. Global *R* is 0 if the similarities of the samples are the same between groups and within groups (Clarke & Warwick, 2001).

To compare the difference in the relative peak area of major categories of compounds among 7 individuals, we computed a Bray–Curtis similarity index using the average of each category of compounds in the ANOSIM test procedure with 1,000 permutations. To examine whether there are correlations between different categories of compounds, we calculated the average of each category of compounds per bat, and then used Pearson correlation to assess the relationship between different categories of compounds.

To assess the effect of sampling date on chemical composition, we performed a NMDS plot of a set of 3 samples from 7 individuals collected over a period of 3 months (day 0, day 45 and day 90, respectively). The NMDS plot was based on Bray–Curtis similarities. To examine whether the chemical composition of each individual changed over time, we compared the chemical composition of gland secretions from 7 individuals at the three sampling dates using the ANOSIM test procedure with 1,000 permutations.

To examine whether the males responded differently towards forehead gland odors from different individuals, we first used Log_10_ transformations to normalize distributions of sniffing duration (Kolmogorove-Smirnov test: *p* > 0.05). For the habituation phase, one-way analysis of variance (ANOVA) was used to test whether there were significant differences in the sniffing duration when the tested bats were repeatedly presented with the habituation odor over 4 days. If there were significant differences, we used Tukey’s multiple-comparison tests to further test the differences in the sniffing duration across days. For the discrimination phase, paired sample *t* tests were used to compare differences in the sniffing durations between the habituation odor and the novel odor.

All individuals used in trials where both bats were capable of gland protrusion and in trials with both bats were incapable of gland protrusion were collected from caves in Yunnan Province. Mixed pairs were derived from two source populations. Sixteen adult males used in the mixed-pair trials were caught in Yunnan Province and 24 adult males were caught in Guizhou Province. This experimental design allowed us to assess the effect of population on contest duration and intensity. The impact of population of origin on contest duration was determined using independent sample *t* tests, and Pearson chi-square tests were used to compare differences in the aggressive displays between bats from Yunnan Province and Guizhou Province.

One-way ANOVA was used to test the effects of vocal behavior on contest duration and frequency of aggressive displays, and Kruskal–Wallis tests were used to assess the impact of vocal behavior on the proportion of aggressive display. In all cases, vocal behavior was treated as present or absent for each member of the pair. Thus vocal behavior was either vocalizations from both contestants, only one contestant or neither contestant.

To test the potential function of the chemical signals in conflict solution, we first used a Kolmogorov-Smirnov test to test the normality of all data. We used parametric tests for contest duration and frequency of aggressive displays, and non-parametric tests for proportion of aggressive displays. One-way ANOVA was used to compare differences in the contest duration and frequency of aggressive displays among the 3 categories of paired bats. If there were significant differences, we used Tukey’s multiple-comparison tests to further test the differences in the contest duration and frequency of aggressive displays between any two categories. Kruskal–Wallis tests were used to examine whether there were differences in the proportion of aggressive displays among the pair categories. Pearson chi-square tests were used to test whether contestants in trials where neither bat had functioning glands tended to be involved in contests with more physical contact than contests where at least one bat had a functioning gland. To assess the effects of the presence of gland protrusion on proportion of winning, we used exact binomial probability tests to estimate whether contests tended to be won by the male that was capable of gland protrusion when the other individual’s gland was glued shut. Paired sample *t* tests were used to compare differences in the frequency of aggressive displays, and Wilcoxon signed–rank tests were used to compare differences in the proportion of aggressive displays between the individual with gland protrusion and the individual with the glued gland.

Statistical analyses of chemical data (except for the Pearson correlation) were performed using VEGAN package in RStudio (version 1.0.136; (R Core Team, 2015). Statistical analyses of behavioral data and Pearson correlation were performed with SPSS 22.0 (IBM Corp., Armonk, NY, U.S.A.). We considered *p* < 0.05 as significant.

## Acknowledgements

This research was supported by the National Natural Science Foundation of China (Grant Nos. 31872680, 31922050, 31670390), the Fund of the Jilin Province Science and Technology Development Project (Grant no. 20180101024JC), and the Program for Introducing Talents to Universities (B16011).

## Author contributions

CMZ, CNS and TLJ designed the study. CMZ, CNS and HG collected the data. CMZ and CNS analyzed the data and wrote the manuscript. TLJ, JF and JRL revised the manuscript. All of the authors contributed critically to the drafts and gave final approval for publication.

## Ethics

All procedures complied with the Guidelines for the Use of Animals in Research (ASAB/ABS, 2017), and the National Natural Science Foundation of China for experiments involving vertebrate animals, and were approved by the Northeast Animal Research Authority of Northeast Normal University, China (approval number: NENU-W-2018-101). There were no visible physical injuries and deaths during the whole experiment period. After completion of the experiments, all the bats were released into their original caves.

## Supplemental materials

**Figure S1.**
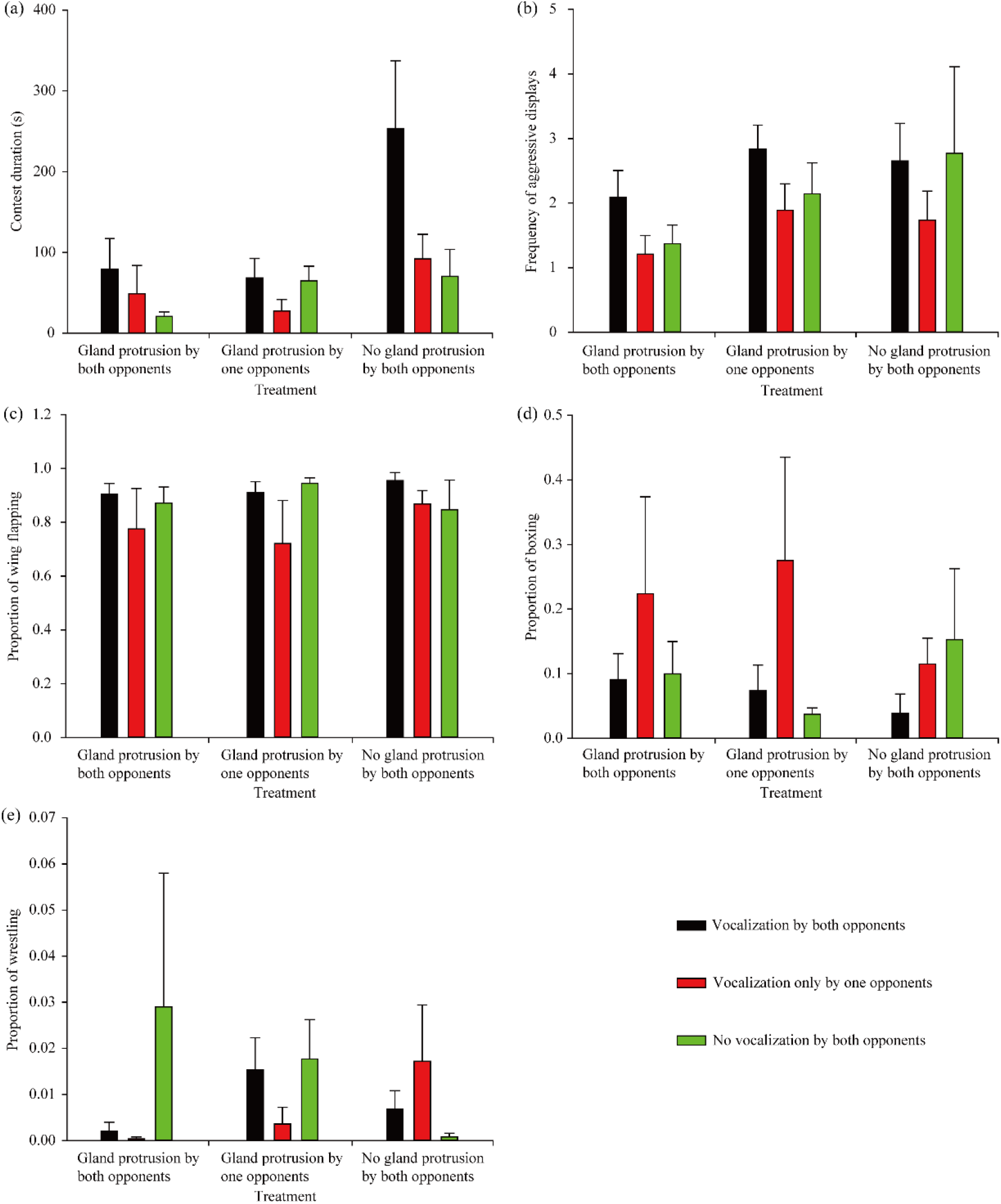
Effect of vocalization on agonistic interactions. Effect of vocalization on contest duration (a), frequency of aggressive displays (b), proportion of wing flapping (c), proportion of boxing (d), and proportion of wrestling (e) among 3 pair types. Black columns = vocalization by both opponents; red column = vocalization only by one opponent; green column = no vocalization by both opponents. Data are shown as means ±standard errors.

**Figure S2.**
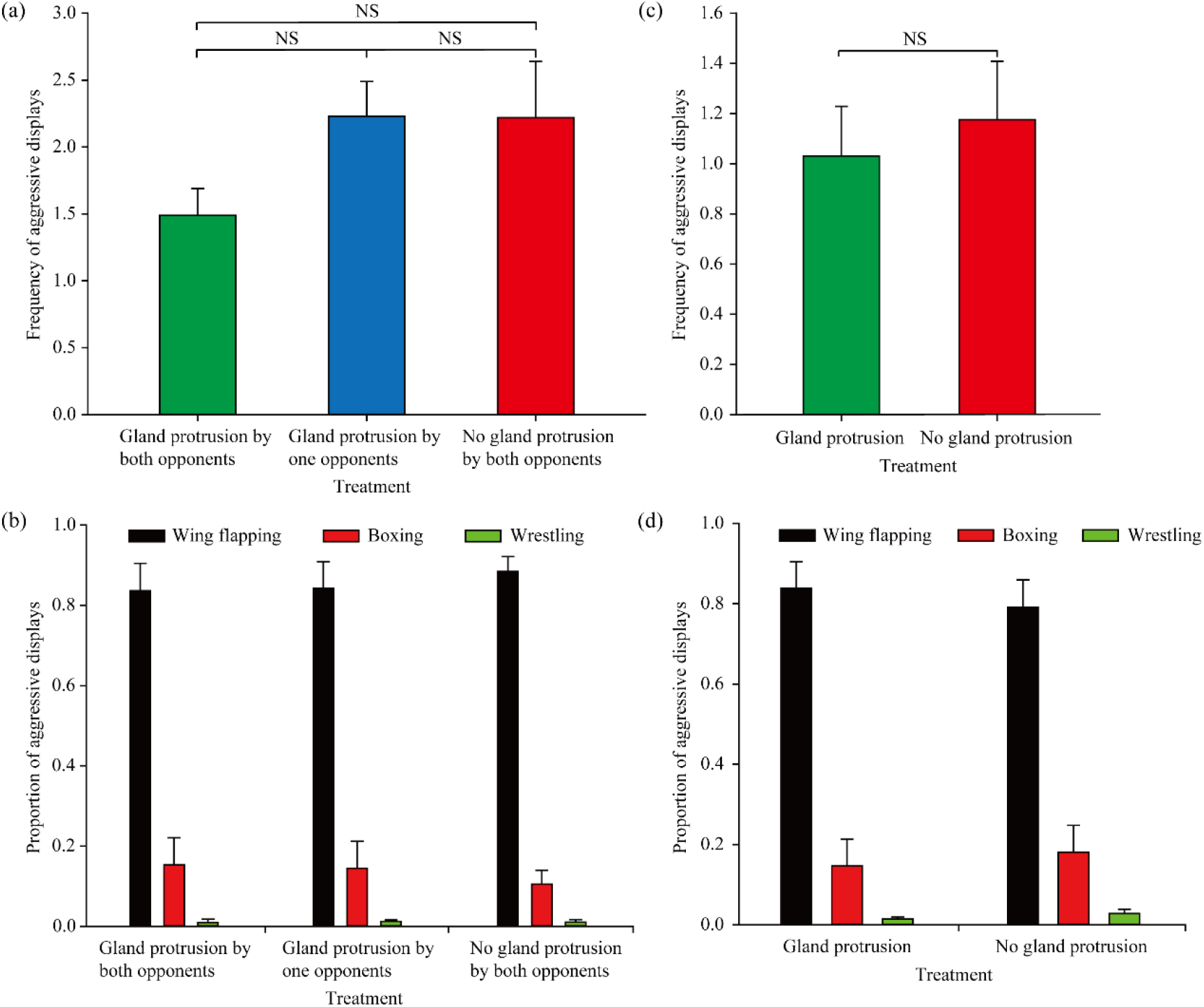
Comparisons of agonistic displays. Comparisons of frequency of aggressive displays (a) and proportion of aggressive displays (b) among 3 pair types. Comparisons of frequency of aggressive displays (c) and proportion of aggressive displays (d) between the individuals with gland protrusion and the individuals with no gland protrusion. Data are shown as means ± standard errors. NS indicate no significant difference (*p* > 0.05).

## References

Adams, D.M., Li, Y. & Wilkinson, G.S. (2018). Male scent gland signals mating status in greater spear-nosed bats, *Phyllostomus hastatus*. J. Chem. Ecol., 44, 975–986. DOI: https://doi.org/10.1007/s10886-018-1003-8

Alberts, A.C. (1992). Constraints on the design of chemical communication systems in terrestrial vertebrates. Am. Nat., 139, S62–S89. DOI: https://doi.org/10.1086/285305

Albone, E.S. (1984). Mammalian semiochemistry: the investigation of chemical signals between mammals. John Wiley & Sons, New York.

Atema, J. & Steinbach, M. (2007). Chemical communication and social behavior of the lobster *Homarus americanus* and other decapod Crustacea. In: Evolutionary ecology of social and sexual systems: crustaceans as model organisms. (ed Duffy, J.E. & Thiel, M.). Oxford University Press, New York, pp. 115–144.

Bastian, A. & Schmidt, S. (2008). Affect cues in vocalizations of the bat, *Megaderma lyra*, during agonistic interactions. J. Acoust. Soc. Am., 124, 598–608. DOI: https://doi.org/10.1121/1.2924123

Bloss, J., Acree, T.E., Bloss, J.M., Hood, W.R. & Kunz, T.H. (2002). Potential use of chemical cues for colony-mate recognition in the big brown bat, *Eptesicus fuscus*. J. Chem. Ecol., 28, 819–834. DOI: https://doi.org/10.1023/A:1015296928423

Bonadonna, F. & Nevitt, G.A. (2004). Partner-specific odor recognition in an Antarctic seabird. Science, 306, 835–835. DOI: https://doi.org/10.1126/science.1103001

Bouchard, S. (2001). Sex discrimination and roostmate recognition by olfactory cues in the African bats, *Mops condylurus* and *Chaerephon pumilus* (Chiroptera: Molossidae). J. Zool., 254, 109–117. DOI: https://doi.org/10.1017/S0952836901000607

Bradbury, J.W. & Vehrencamp, S.L. (2011). Principles of animal communication. 2nd ed. Sinauer Associates, Sunderland, MA.

Breithaupt, T. & Atema, J. (2000). The timing of chemical signaling with urine in dominance fights of male lobsters *(Homarus americanus)*. Behav. Ecol. Sociobiol., 49, 67–78. DOI: https://doi.org/10.1007/s002650000271

Breithaupt, T. & Eger, P. (2002). Urine makes the difference: chemical communication in fighting crayfish made visible. J. Exp. Biol., 205, 1221–1231. DOI: https://doi.org/10.1086/315276

Briffa, M. (2015). Agonistic signals: integrating analysis of functions and mechanisms. In: Animal signaling and function, an integrative approach (ed Irschick, D.J., Briffa, M. & Podos, J.). Wiley Blackwell, Hoboken, NJ, pp. 141–167.

Briffa, M. & Elwood, R.W. (2004). Use of energy reserves in fighting hermit crabs. Proc. R. Soc. Lond. B, 271, 373–379. DOI: http://doi.org/10.1098/rspb.2003.2633

Briffa, M. & Sneddon, L.U. (2007). Physiological constraints on contest behaviour. Funct. Ecol., 21, 627–637. DOI: https://doi.org/10.1111/j.1365-2435.2006.01188.x

Brooke, A.P. & Decker, D.M. (1996). Lipid compounds in secretions of fishing bat, *Noctilio leporinus* (Chiroptera: Noctilionidae). J. Chem. Ecol., 22, 1411–1428. DOI: https://doi.org/10.1007/BF02027721

Burgener, N., Dehnhard, M., Hofer, H. & East, M.L. (2009). Does anal gland scent signal identity in the spotted hyaena? Anim. Behav., 77, 707–715. DOI: https://doi.org/10.1016/j.anbehav.2008.11.022

Bushdid, C., Magnasco, M.O., Vosshall, L.B. & Keller, A. (2014). Humans can discriminate more than 1 trillion olfactory stimuli. Science, 343, 1370–1372. DOI: https://doi.org/10.1126/science.1249168

Caspers, B., Franke, S. & Voigt, C.C. (2008). The wing-sac odour of male greater sac-winged bats *Saccopteryx bilineata* (Emballonuridae) as a composite trait: seasonal and individual differences. In: Chemical signals in vertebrates 11 (ed Hurst, J.L., Beynon, R. J., Roberts, S.C. & Wyatt, T.D.). Springer, New York, pp. 151–160.

Caspers, B.A., Schroeder, F.C., Franke, S., Streich, W.J. & Voigt, C.C. (2009). Odour-based species recognition in two sympatric species of sac-winged bats (*Saccopteryx bilineata, S. leptura*): combining chemical analyses, behavioural observations and odour preference tests. Behav. Ecol. Sociobiol., 63, 741–749. DOI: https://doi.org/10.1007/s00265-009-0708-7

Cheng, H.C. & Lee, L.L. (2004). Temporal variations in the size and composition of Formosan leaf-nosed bat (*Hipposideros terasensis*) colonies in central Taiwan. Zool. Stud., 43, 787–794.

Chouinard, A.J., Wilburn, D.B., Houck, L.D. & Feldhoff, R.C. (2013). Individual variation in pheromone isoform ratios of the red-legged salamander, *Plethodon shermani*. In: Chemical signals in vertebrates 12. Springer, pp. 99–115.

Clement, M., Dietz, N., Gupta, P. & Kanwal, J. (2006). Audiovocal communication and social behavior in mustached bats. In: Behavior and neurodynamics for auditory communication (ed Kanwal, J.S. & Ehret, G.). Cambridge University Press, Cambridge, pp. 57–84.

Drea, C.M., Boulet, M., Delbarco-Trillo, J., Greene, L.K., Sacha, C.R., Goodwin, T.E. & Dubay, G.R. (2013). The ‘secret’ in secretions: methodological considerations in deciphering primate olfactory communication. Am. J. Primatol., 75, 621–642. DOI: https://doi.org/10.1002/ajp.22143

Enquist, M. & Leimar, O. (1990). The evolution of fatal fighting. Anim. Behav., 39, 1–9. DOI: https://doi.org/10.1016/S0003-3472(05)80721-3

Faulkes, C.G., Elmore, J.S., Baines, D.A., Fenton, B., Simmons, N.B. & Clare, E.L. (2019). Chemical characterisation of potential pheromones from the shoulder gland of the Northern yellow-shouldered-bat, *Sturnira parvidens* (Phyllostomidae: Stenodermatinae). Peer J, 7, e7734. DOI: https://doi.org/10.7717/peerj.7734

Fernandez, A.A., Fasel, N., Knörnschild, M. & Richner, H. (2014). When bats are boxing: aggressive behaviour and communication in male Seba’s short-tailed fruit bat. Anim. Behav., 98, 149–156. DOI: https://doi.org/10.1016/j.anbehav.2014.10.011

Green, P. & Patek, S. (2015). Contests with deadly weapons: telson sparring in mantis shrimp (Stomatopoda). Biol. Lett., 11, 20150558. DOI: https://doi.org/10.1098/rsbl.2015.0558

Green, P. & Patek, S. (2018). Mutual assessment during ritualized fighting in mantis shrimp (Stomatopoda). Proc. R. Soc. B, 285, 20172542. DOI: https://doi.org/10.1098/rspb.2017.2542

Gustin, M.K. & McCracken, G.F. (1987). Scent recognition between females and pups in the bat *Tadarida brasiliensis mexicana*. Anim. Behav., 35, 13–19. DOI: https://doi.org/10.1016/S0003-3472(87)80205-1

Hagey, L. & MacDonald, E. (2003). Chemical cues identify gender and individuality in giant pandas *(Ailuropoda melanoleuca)*. J. Chem. Ecol., 29, 1479–1488. DOI: https://doi.org/10.1023/A:1024225806263

Halpin, Z.T. (1986). Individual odors among mammals: origins and functions. Adv. Stud. Behav., 16, 39–70. DOI: https://doi.org/10.1016/S0065-3454(08)60187-4

Hurst, J.L., Payne, C.E., Nevison, C.M., Marie, A.D., Humphries, R.E., Robertson, D.H., Cavaggioni, A. & Beynon, R.J. (2001). Individual recognition in mice mediated by major urinary proteins. Nature, 414, 631–634. DOI: https://doi.org/10.1038/414631a

Kind, T., Wohlgemuth, G., Lee, D.Y., Lu, Y., Palazoglu, M., Shahbaz, S. & Fiehn, O. (2009). FiehnLib: mass spectral and retention index libraries for metabolomics based on quadrupole and time-of-flight gas chromatography/mass spectrometry. Anal. Chem., 81, 10038–10048. DOI: https://doi.org/10.1021/ac9019522

Lenoir, A., Fresneau, D., Errard, C. & Hefetz, A. (1999). Individuality and colonial identity in ants: The emergence of the social representation concept. In: Information Processing in Social Insects (ed Detrain, C., Deneubourg, J.L. & Pasteels, J.M.). Birkhäuser Verlag, Basel, Switzerland, pp. 219–237.

Liebal, K., Waller, B.M., Burrows, A.M. & Slocombe, K.E. (2014). Primate communication: a multimodal approach. Cambridge University Press, Cambridge.

Logue, D., Abiola, I., Rains, D., Bailey, N., Zuk, M. & Cade, W. (2010). Does signalling mitigate the cost of agonistic interactions? A test in a cricket that has lost its song. Proc. R. Soc. B, 277, 2571–2575. DOI: https://doi.org/10.1098/rspb.2010.0421

Mardon, J., Saunders, S.M., Anderson, M.J., Couchoux, C. & Bonadonna, F. (2010). Species, gender, and identity: cracking petrels’ sociochemical code. Chem. senses, 35, 309–321. DOI: https://doi.org/10.1093/chemse/bjq021

Mateo, J.M. (2006). The nature and representation of individual recognition odours in Belding’s ground squirrels. Anim. Behav., 71, 141–154. DOI: https://doi.org/10.1016/j.anbehav.2005.04.006

Moore, P.J., Reagan-Wallin, N.L., Haynes, K.F. & Moore, A.J. (1997). Odour conveys status on cockroaches. Nature, 389, 25–25. DOI: https://doi.org/10.1038/37888

Moreira, P.L., López, P. & Martín, J. (2006). Femoral secretions and copulatory plugs convey chemical information about male identity and dominance status in Iberian rock lizards *(Lacerta monticola)*. Behav. Ecol. Sociobiol., 60, 166. DOI: https://doi.org/10.1007/s00265-005-0153-1

Novotny, M., Harvey, S., Jemiolo, B. & Alberts, J. (1985). Synthetic pheromones that promote inter-male aggression in mice. Proc. Natl. Acad. Sci. USA, 82, 2059–2061. DOI: https://doi.org/10.1073/pnas.82.7.2059

R Development Core Team. (2015). R: a language and environment for statistical computing. Vienna: R Foundation for Statistical Computing. Available at http://www.R-project.org/.

Rehorek, S.J., Smith, T.D. & Bhatnagar, K.P. (2010). The orbitofacial glands of bats: An investigation of the potential correlation of gland structure with social organization. Anat. Rec., 293, 1433–1448. DOI: https://doi.org/10.1002/ar.21046

Rendon, N.M., Soini, H.A., Scotti, M.-A.L., Novotny, M.V. & Demas, G.E. (2016). Urinary volatile compounds differ across reproductive phenotypes and following aggression in male Siberian hamsters. Physiol. Behav., 164, 58–67. DOI: https://doi.org/10.1016/j.physbeh.2016.05.034

Safi, K. & Kerth, G. (2003). Secretions of the interaural gland contain information about individuality and colony membership in the Bechstein’s bat. Anim. Behav., 65, 363–369. DOI: https://doi.org/10.1006/anbe.2003.2067

Schultze-Westrum, T.G. (1969). Social communication by chemical signals in flying phalangers (Petaurus breviceps papuanus). In: Olfaction and taste: proceedings of the third international symposium (ed Pfaffman, C.). Rockefeller University Press, New York, pp. 268–277.

Scully, W.M., Fenton, M. & Saleuddin, A.S. (2000). A histological examination of the holding sacs and glandular scent organs of some bat species (Emballonuridae, Hipposideridae, Phyllostomidae, Vespertilionidae, and Molossidae). Can. J. Zool., 78, 613–623. DOI: https://doi.org/10.1139/z99-248

Smith, T.E., Tomlinson, A.J., Mlotkiewicz, J.A. & Abbott, D.H. (2001). Female marmoset monkeys *(Callithrix jacchus)* can be identified from the chemical composition of their scent marks. Chem. Senses, 26, 449–458. DOI: https://doi.org/10.1093/chemse/26.5.449

Sorensen, P.W. (2015). Introduction to pheromones and related cues in fishes. In: Fish pheromones and related cues (ed Sorensen, P.W. & Wisenden, B.D.). John Wiley & Sons, Iowa, USA, pp. 1–9.

Stritih, N. & Kosi, A.Ž. (2017). Olfactory signaling of aggressive intent in male-male contests of cave crickets (*Troglophilus neglectus*; Orthoptera: Rhaphidophoridae). PLoS ONE, 12, e0187512. DOI: https://doi.org/10.1371/journal.pone.0187512

Sun, C., Jiang, T., Kanwal, J.S., Guo, X., Luo, B., Lin, A. & Feng, J. (2018). Great Himalayan leaf-nosed bats modify vocalizations to communicate threat escalation during agonistic interactions. Behav. Process., 157, 180–187. DOI: https://doi.org/10.1016/j.beproc.2018.09.013

Sun, C., Zhang, C., Gu, H., Jiang, T. & Feng, J. (2019). Self-assessment strategy during contest decisions between male Great Himalayan leaf-nosed bats. Behav. Ecol. Sociobiol., 73, 45. DOI: https://doi.org/10.1007/s00265-019-2657-0

Tibbetts, E.A. & Dale, J. (2007). Individual recognition: it is good to be different. Trends Ecol. Evol., 22, 529–537. DOI: https://doi.org/10.1016/j.tree.2007.09.001

Tibbetts, E.A. & Lindsay, R. (2008). Visual signals of status and rival assessment in Polistes dominulus paper wasps. Biol. Lett., 4, 237–239. DOI: https://doi.org/10.1098/rsbl.2008.0048

Voigt, C.C., Behr, O., Caspers, B., von Helversen, O., Knörnschild, M., Mayer, F. & Nagy, M. (2008). Songs, scents, and senses: sexual selection in the greater sac-winged bat, *Saccopteryx bilineata*. J. Mammal., 89, 1401–1410. DOI: https://doi.org/10.1644/08-MAMM-S-060.1

Wyatt, T.D. (2014). Pheromones and animal behavior: chemical signals and signatures. Cambridge University Press, Cambridge.

Yang, Y.-J. (2011). Mating system and kinship of the formasan leaf-nosed bat, *Hipposideros armiger Terasensis* (Chiroptera, Hippsideridae). Dissertation. National Chung Hsing University.

Zhao, X., Jiang, T., Gu, H., Liu, H., Sun, C., Liu, Y. & Feng, J. (2018). Are aggressive vocalizations the honest signals of body size and quality in female Asian particoloured bats? Behav. Ecol. Sociobiol., 72, 96. DOI: https://doi.org/10.1007/s00265-018-2510-x

