## Supplemental Files 1 for "The potential function of forehead gland secretions in conflict resolution of the male Great Himalayan leaf-nosed bats": Supplemental materials.docx

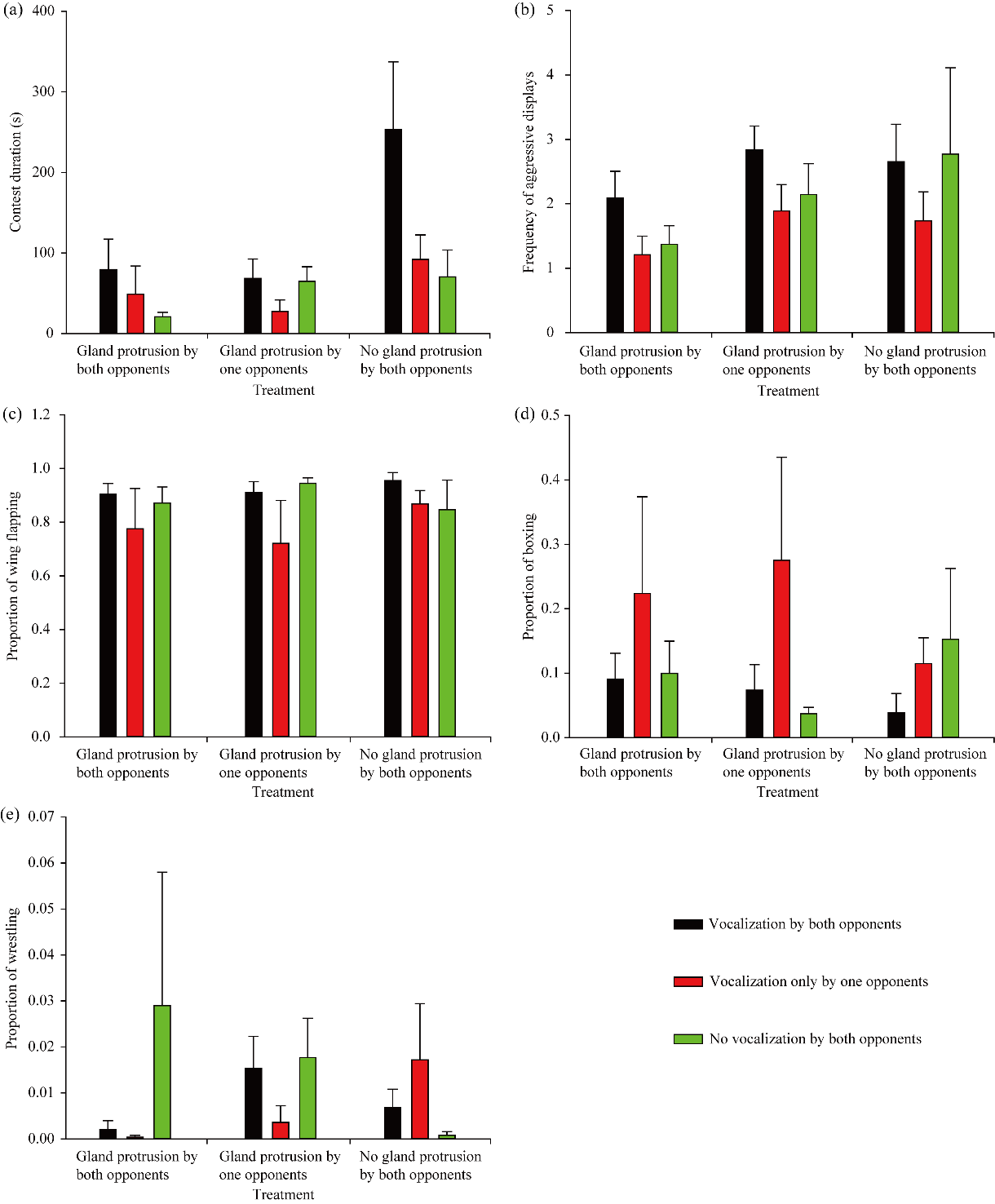

**Figure S1.** Effect of vocalization on agonistic interactions. Effect of vocalization on contest duration (a), frequency of aggressive displays (b), proportion of wing flapping (c), proportion of boxing (d), and proportion of wrestling (e) among 3 pair types. Black columns = vocalization by both opponents; red column = vocalization only by one opponent; green column = no vocalization by both opponents. Data are shown as means ± standard errors.

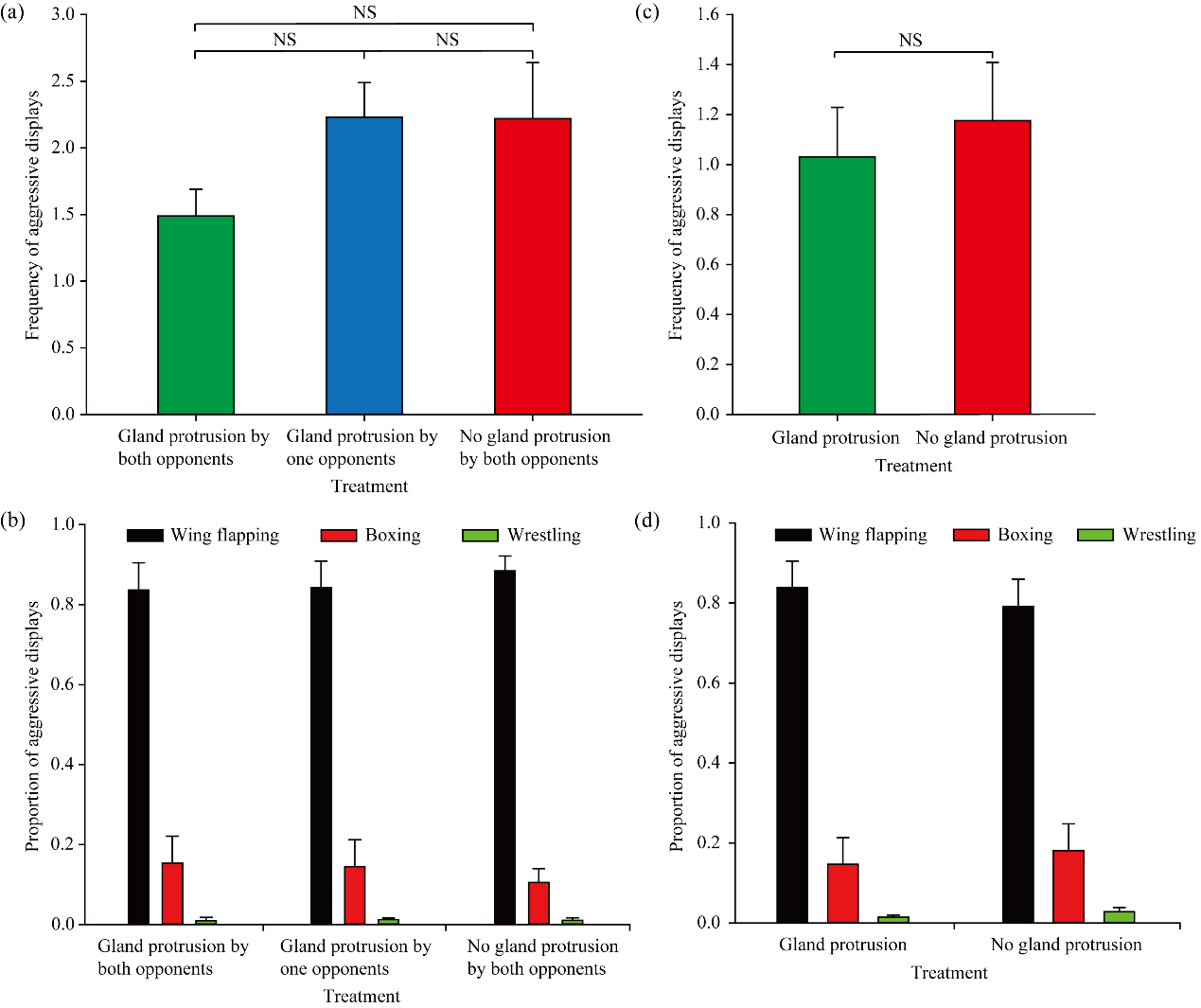

**Figure S2.** Comparisons of agonistic displays. Comparisons of frequency of aggressive displays (a) and proportion of aggressive displays (b) among 3 pair types. Comparisons of frequency of aggressive displays (c) and proportion of aggressive displays (d) between the individuals with gland protrusion and the individuals with no gland protrusion. Data are shown as means ± standard errors. NS indicate no significant difference (*p* > 0.05).
