## Supplemental Files 1 for "The potential function of forehead gland secretions in conflict resolution of the male Great Himalayan leaf-nosed bats": Video Legend.docx

**Supplemental materials**

**Video S1.** Video of male *Hipposideros armiger* during agonistic interactions with gland protrusion manipulation. Filmed with two infrared high-speed cameras at 85 frames/s and played back at 25 frames/s.
